# Global Mapping of Mouse CSF Flow via HEAP-METRIC Phase-contrast MRI

**DOI:** 10.1101/2021.09.04.459006

**Authors:** Juchen Li, Mengchao Pei, Binshi Bo, Xinxin Zhao, Jing Cang, Fang Fang, Zhifeng Liang

## Abstract

Roles of Cerebrospinal fluid (CSF) in brain waste clearance and homeostasis has been increasingly recognized, thus measuring its flow dynamics could provide important information about its function and perturbance. While phase-contrast MRI can be used for non-invasive flow mapping, so far its mapping of low velocity flow (such as mouse brain CSF) is not possible. Here we developed a novel generalized Hadamard encoding based multi-band acceleration scheme dubbed HEAP-METRIC (<u>H</u>adamard <u>E</u>ncoding <u>AP</u>proach of <u>M</u>ulti-band <u>E</u>xcitation for short <u>TR</u> <u>I</u>maging a<u>C</u>celerating), and with significantly increased SNR per time, HEAP-METRIC phase-contrast MRI achieved fast and accurate mapping of slow (~10^2^ micron/s) flow. Utilizing this novel method, we revealed a heterogeneous global pattern of CSF flow in the awake mouse brain with a averaged flow of ~200 micron/s, and further found isoflurane anesthesia reduced CSF flow that was accompanied by reduction of glymphatic function. Therefore, we developed the novel HEAP-METRIC phase-contrast MRI for mapping low velocity flow, and demonstrated its capability for global mapping of mouse CSF flow and its potential alterations related to various physiological or pathological conditions.

## Introduction

Cerebrospinal fluid (CSF) has been increasingly recognized as an important component in brain waste clearance and homeostasis. Produced by choroid plexus in ventricles, CSF is traditionally believed to flow from the ventricular system into the subarachnoid space and exit via cranial and spinal nerves and arachnoid granulations^1^. Recent proposed glymphatic system theory^2^ further extends the role of CSF, as it flows through the brain parenchyma along the paravascular space and exchanges with interstitial fluid (ISF). The (dys)function of glymphatic system has been implicated in various physiological and pathological processes, such as sleep, aging and neurodegenerative diseases^3,4^.

As an integral part of brain’s waste clearance system, the CSF flow dynamics might play an important role in this process. Studies found that ageing related reduction of the CSF flow velocity and its association with cognitive deficits^5,6^. Moreover, the CSF flow is positively correlated with the CSF protein level, suggesting its role in protein clearance from the brain^7^.

However, the CSF flow measurement in rodents has been difficult: invasive methods would most likely perturb the flow, while non-invasive methods such as conventional phase-contrast MRI (PC-MRI) cannot map the presumably low velocity CSF flow in the rodent brain. Phase contrast MR pulse sequence is widely used for flow velocity quantification (e.g., blood or CSF flow in human). However, human CSF flow is on the order of ~10^4^ μm/s^8,9^ and with much small brain size in rodents, CSF flow in rodents would likely be several orders of magnitude lower. Such extremely slow flow cannot be readily mapped using conventional PC-MRI: firstly, low SNR phase maps are heavily contaminated by Rician noise and thus create large bias errors in velocity images^10–12^; Secondly, such slow flow requires high amplitude of toggling gradients in PC-MRI, which leads to high eddy current field induced phase errors^13–15^. Two factors combined, mapping low velocity flow such as CSF in rodents has not been achieved.

Therefore, we developed a novel PC-MRI method based on a generalized Hadamard multi-band (MB) acceleration approach dubbed HEAP-METRIC (<u>H</u>adamard <u>E</u>ncoding <u>AP</u>proach of <u>M</u>ulti-band <u>E</u>xcitation for short <u>TR</u> <u>I</u>maging a<u>C</u>celerating) PC-MRI, which can readily achieve a MB band of ~20. We applied this novel method to achieve global mapping of mouse ventricular CSF flow for the first time, and found a spatially heterogeneous CSF flow pattern with an averaged flow around 216 μm/s in the awake mouse brain. Importantly, we found a reduction of CSF flow in the anesthetized condition compared to the awake condition, further highlighting HEAP-METRIC PC-MRI’s ability to detect physiological or pathological alterations of CSF flow in rodents. HEAP-METRIC PC-MRI can be openly accessed (https://github.com/ZhifengLiangLab/HEAP-METRIC).

## Results

### Design of HEAP-METRIC PC-MRI and its validation for low velocity mapping

SNR may be significantly improved by Hadamard multi-band (MB) approach in multi-slice fast imaging with short repetition time (TR), as it enables multiple signal averages without extra scanning time. Previous Hadamard MB Encoding methods^16–18^ used phase modulated Radiofrequency (RF) waveforms to simultaneously excite and acquire multi-slice signals, thus accelerating the imaging acquisition. In the current work, we developed a generalized Hadamard encoding scheme using complex Hadamard matrix (see Methods) which is dubbed HEAP-METRIC (<u>H</u>adamard <u>E</u>ncoding <u>AP</u>proach of <u>M</u>ulti-band <u>E</u>xcitation for short <u>TR</u> <u>I</u>maging a<u>C</u>celerating), allowing MB factor to be any arbitrary number. Fig. 1 illustrates the approach we developed (see Methods for details): with Hadamard encoding phase modulation a series of RF waveforms W_n_(t) were designed (Fig. 1a); and the MB excitation profiles M_n_(ω) were shown in Fig. 1b, with each band signal phase encoded in complex domain based on Hadamard matrix. The above generalized Hadamard encoding was applied in mouse CSF mapping (Fig. 1c). After Hadamard decoding reconstruction (Fig. 1d), phase unwrapping, eddy current field shimming (Fig. 1e) and combining three directional results yielded quantitative CSF velocity maps (Fig. 1f).

**Figure 1.**
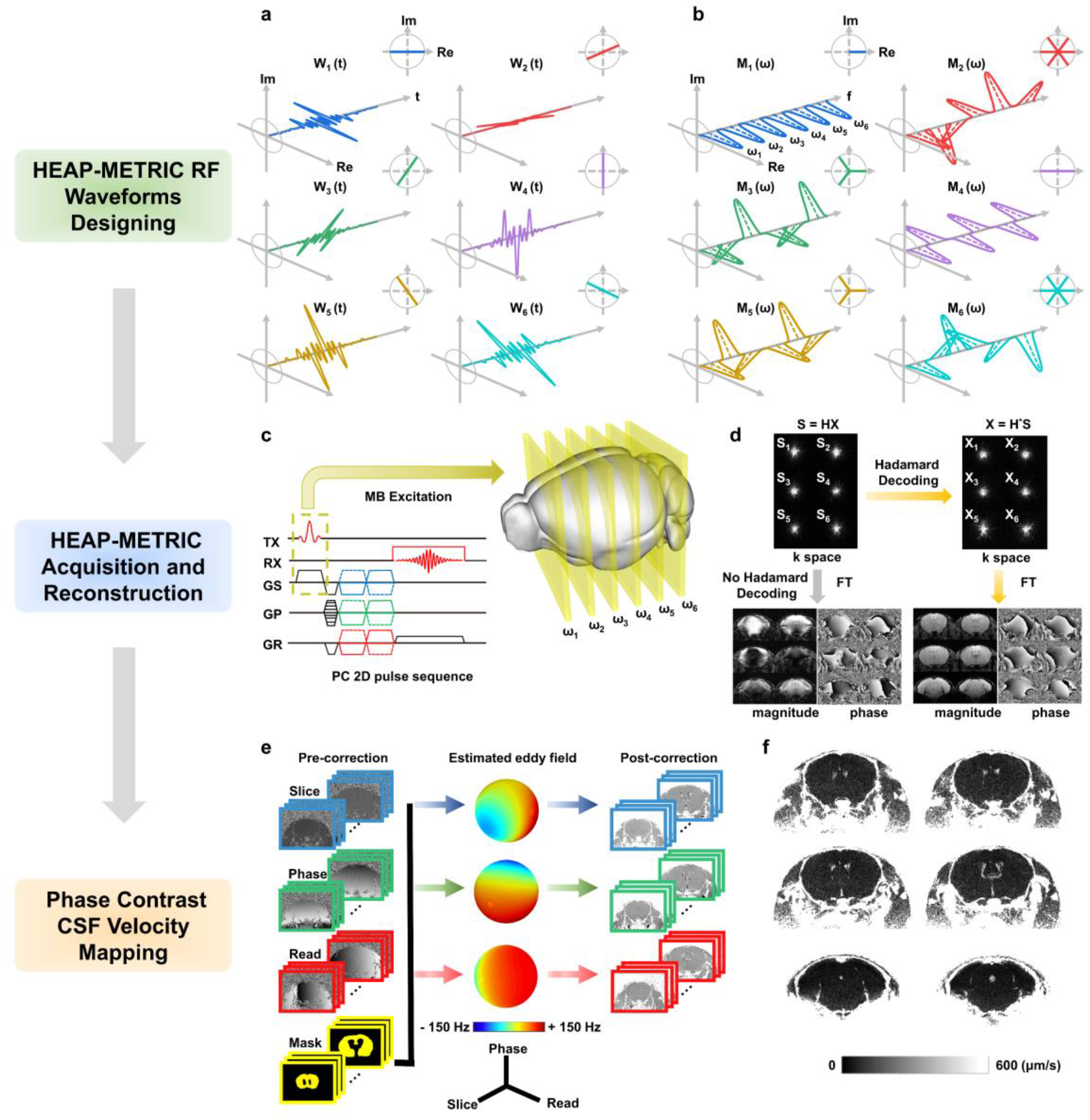
HEAP-METRIC PC-MRI for mouse CSF velocity mapping. a, HEAP-METRIC RF waveforms using MB factor 6 as an example (W_1_(t)-W_6_(t)). Each waveform was plotted as a time dependent 3D complex pulse with a projection on complex plane at its upper right corner. Note in actual experiment MB factor 18 was used. b, excitation profiles M_1_(ω)- M_6_(ω) of W_1_(t)-W_6_(t) calculated by Bloch equations. Each profile was plotted as a frequency dependent 3D complex curve with a projection on a complex plane at its upper right corner. Note that each band of each profile rotated on the complex plan by a certain phase angle according to Hadamard matrix. c, illustration of the designed six-band HEAP-METRIC RF pulses applied on a PC 2D sequence for mouse CSF velocity mapping. d, reconstruction of HEAP-METRIC data. The k-space data were Hadamard decoded and Fourier transformed to obtain magnitude and phase images. Note images reconstructed without Hadamard decoding were also displayed in lower left as a comparison but not used. e, eddy current field shimming. Left column, raw phase contrast images of three velocity encoding directions (slice, phase and read) and manually drawn mask on static tissues. Middle column, three spherical maps of eddy current field estimated by 2^nd^-order spatial polynomial fitting on the static field. Right column, phase images after phase correction including unwrapping and shimming. f, final quantitative CSF velocity maps after combination of three directional results.

The substantial SNR gain from HEAP-METRIC acceleration is critical for mapping low velocity flow (Fig. 2). Without any acceleration, conventional PC-MRI could not yield reasonable CSF flow in mouse brain (Fig. 2, single band and low MB factors). Higher MB factors, enabled by HEAP-METRIC, clearly lead to increasingly smaller bias error reduction in velocity maps (Fig. 2a, b). Importantly, our HEAP-METRIC PC-MRI was validated using a slow flow phantom (Fig. 3), as the mapping results showed very high correlation with set averaged flow velocities ranging from 100-600 μm/s. Therefore, HEAP-METRIC PC-MRI clearly demonstrated fast and accurate multi-slice mapping of low velocity flow, and can be openly accessed (https://github.com/ZhifengLiangLab/HEAP-METRIC).

**Figure 2.**
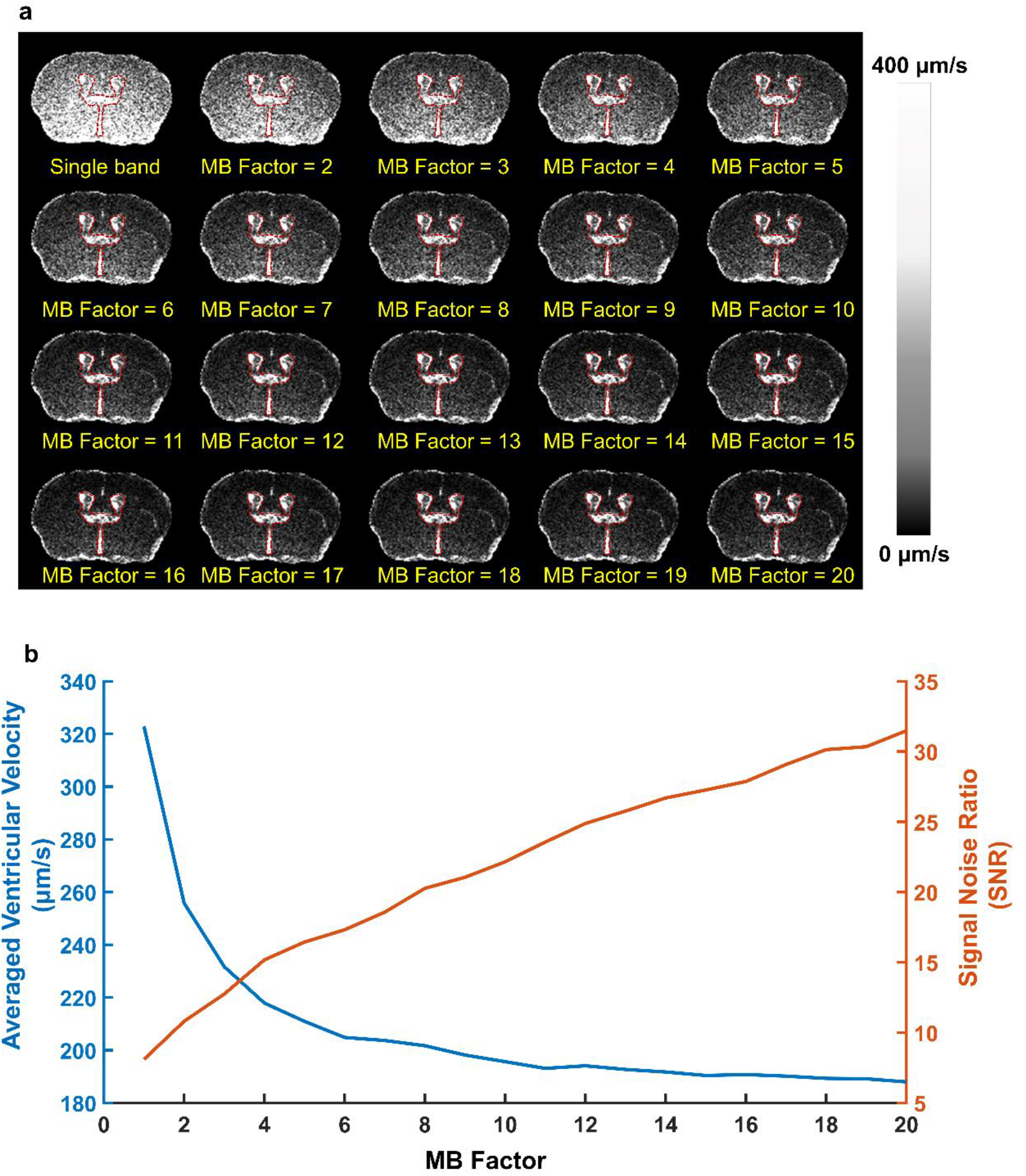
Reduction of bias errors in velocity mapping with SNR improvement. a, reconstructed quantitative velocity maps from conventional single band to MB factors from 2-20 showed notable reduction of bias errors with increasing MB factors. Red outlines, manually drawn CSF ventricle ROI. b, averaged ventricular velocity and SNR (based on CSF ROI in a) exhibited decreasing and increasing trends with increasing MB factors, respectively, which clearly showed high SNR provided by HEAP-METRIC scheme was critical for accurate measurement of low velocity CSF flow in mouse brain.

**Figure 3.**
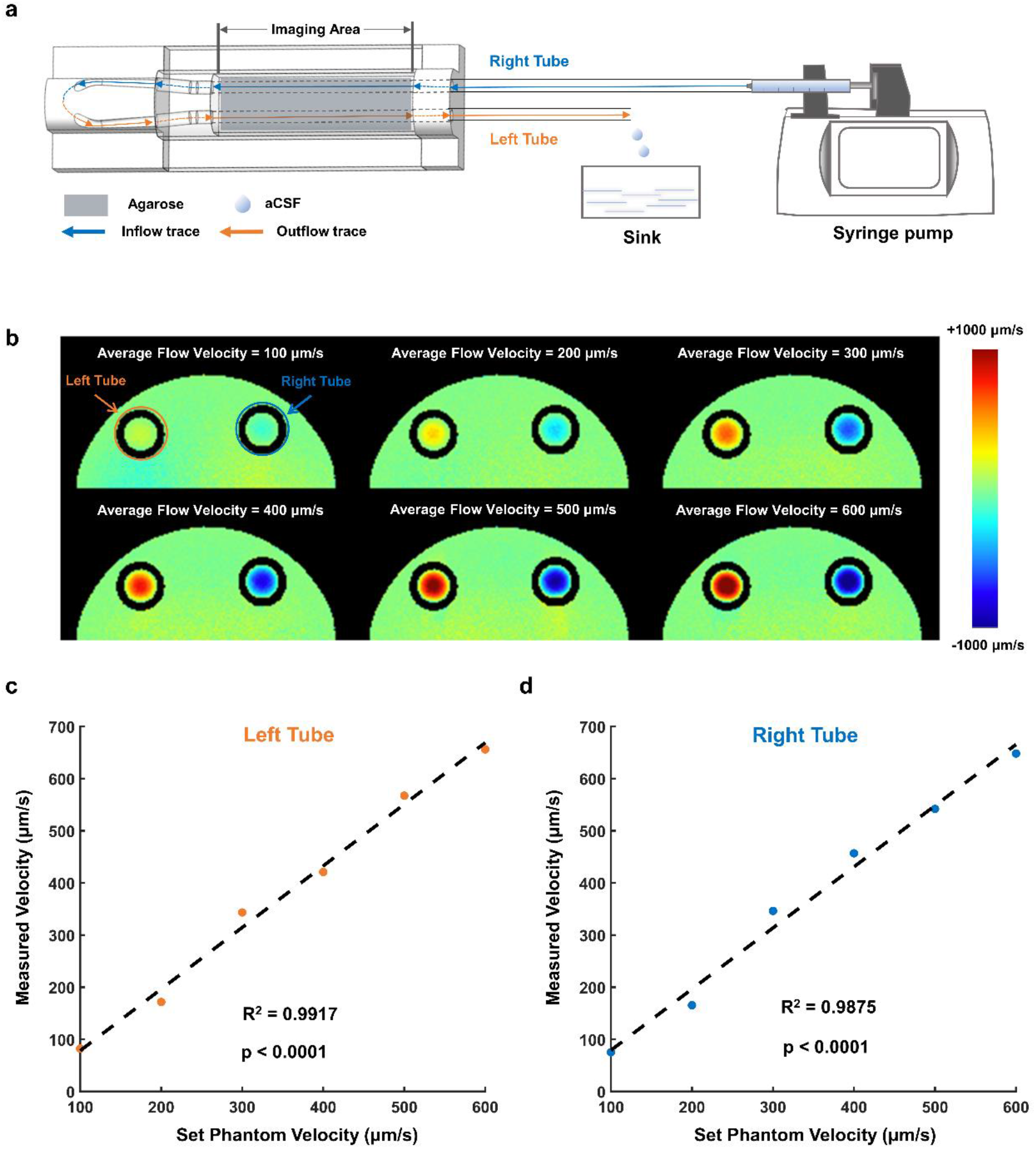
Phantom experiment validates HEAP-METRIC PC-MRI velocity mapping. a, set up of slow flow phantom. b, HEAP-METRIC velocity mapping of flow phantom with set averaged flow velocity from 100-600 μm/s. c and d, high correlation between the measured velocity and the set phantom velocity in the left and right tube, respectively.

### Global mapping of mouse ventricular CSF flow utilizing HEAP-METRIC PC-MRI

Equipped with this novel method and combined with our previously established awake mouse MRI method^19,20^, we systematically characterized ventricular CSF flow at the whole brain level in the awake mice for the first time, and further revealed the impact of anesthesia on CSF flow. With pilot data (Fig. 2) and considering the mouse brain size, we used MB factor of 18 to achieve global CSF mapping (Fig. 4). High resolution (0.08×0.08mm in plane resolution and 36 0.4mm slices) PC-MRI was completed in 21.8 mins with MB factor of 18. Without HEAP-METRIC acceleration, conventional single band acquisition would require 388.8 mins and be clearly infeasible. Globally, CSF flowed at very low velocity (average velocity 216.89 μm/s, Fig. 4e) in the awake condition and exhibited spatial heterogeneity (Fig. 4b, e). To the best of our knowledge, this is the first report of CSF flow mapping in the mouse brain. Not surprisingly, CSF flowed at higher speeds in narrow space such as cerebral aqueduct compared to much larger lateral ventricle (Fig. 4b, e). In addition to the velocity, we observed clear directions of CSF flow in various structures including lateral ventricle, third ventricle and cerebral aqueduct (Fig. 5), which generally agrees with the current understanding of CSF flow directions. For example, the direction of CSF flow in the lateral ventricle was mainly dorsal-ventral, while in the third ventricle and cerebral aqueduct, it was mainly anterior-posterior (slice direction).

**Figure 4.**
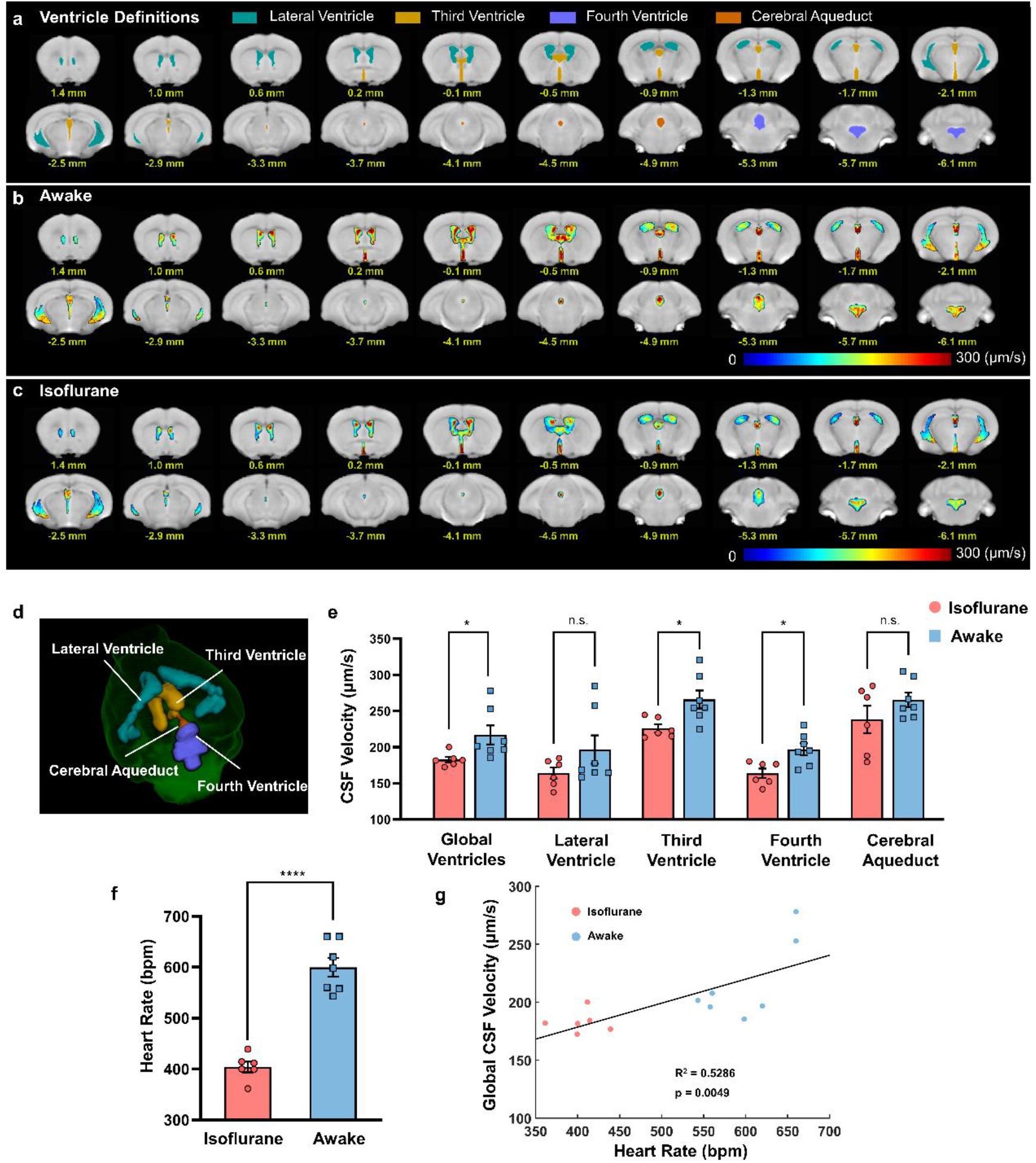
Global mapping of mouse ventricular CSF flow in awake condition and its reduction by isoflurane anesthesia. a, illustration of atlas-based cerebral ventricular definition. b and c, cerebral ventricular CSF velocity mapping in the awake (b, n=6) and isoflurane anesthesia (c, n=7) conditions. Numbers under each slice denoted relative distance from bregma. d, illustration of 3D reconstructed cerebral ventricles. e, reduction of CSF flow by isoflurane anesthesia at global and individual ventricle level. global ventricles: p = 0.0406, third ventricle: p = 0.0184, fourth ventricle: p = 0.01. f, reduction of heart rate in isoflurane anesthesia compared to the awake condition. ****, p<0.0001. g, significant correlation between heart and global CSF velocity.

**Figure 5.**
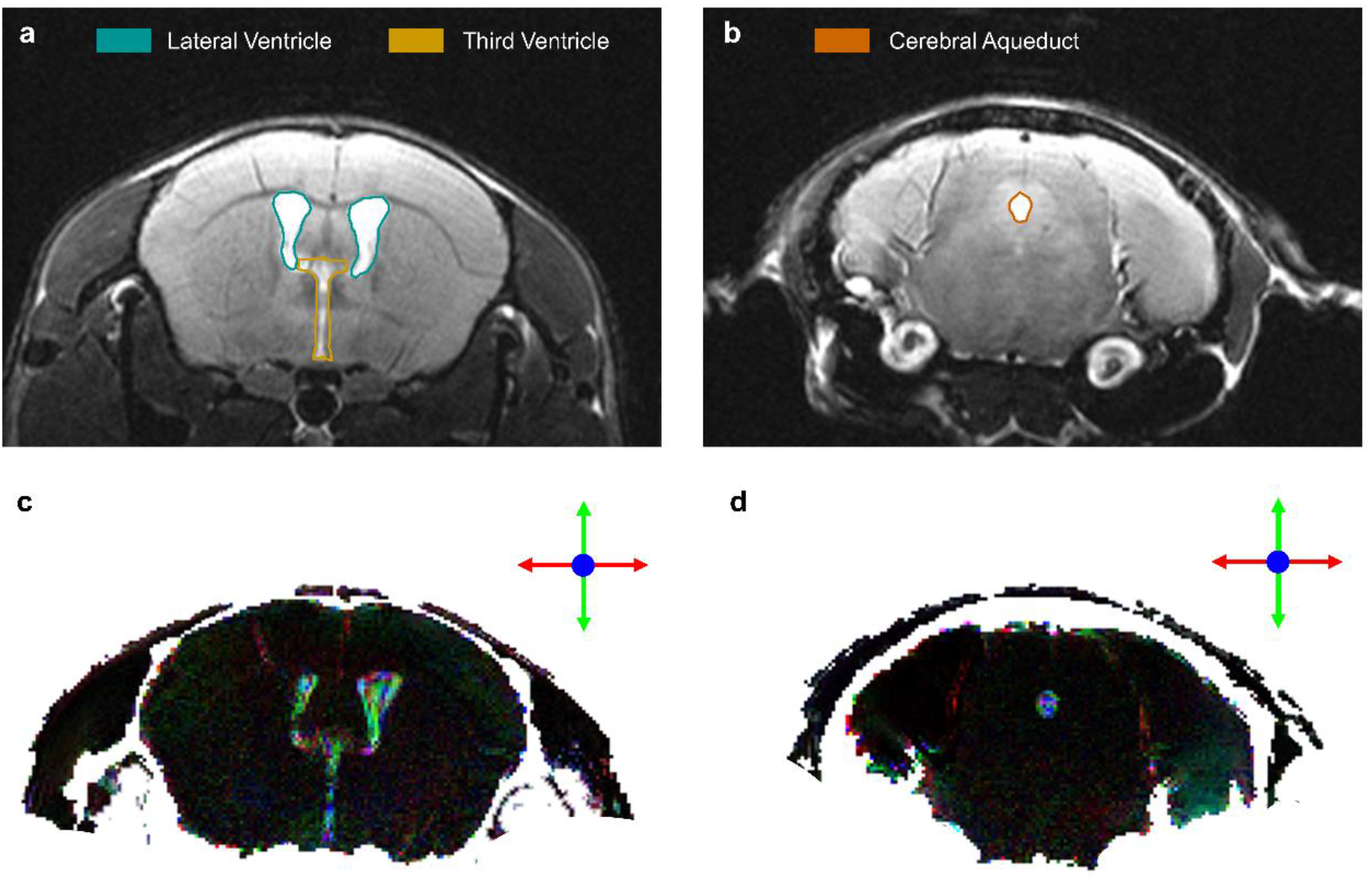
Representative CSF flow directions revealed by HEAP-METRIC mapping. a and b, representative anatomical MRI slices showing lateral and third ventricles, and cerebral aqueduct. c and d, representative color vector maps in the same slices showing directional CSF flow. Green, red and blue color represent vertical (dorsal-ventral), horizontal (medial-lateral) and slice (anterior-posterior) direction, respectively.

### Isoflurane anesthesia reduced CSF velocity and glymphatic function

Importantly, HEAP-METRIC PC-MRI revealed significant reduction of CSF velocity under isoflurane anesthesia (Fig. 4c, e), which has not been reported before. Such reduction was not spatially uniform, as it was more pronounced in third and fourth ventricles. Furthermore, we found significant correlation between heart rate and averaged CSF velocity (Fig. 4f, g), suggesting a possible physiological basis for anesthesia induced CSF velocity reduction. Interestingly, the same isoflurane anesthesia was found to reduce the glymphatic function (as measured by dynamic contrast enhanced MRI, DCE-MRI) compared to the awake condition (Fig. 6), suggesting a potential link between macroscopic ventricular CSF flow and microscopic CSF-ISF flow in brain parenchyma.

**Figure 6.**
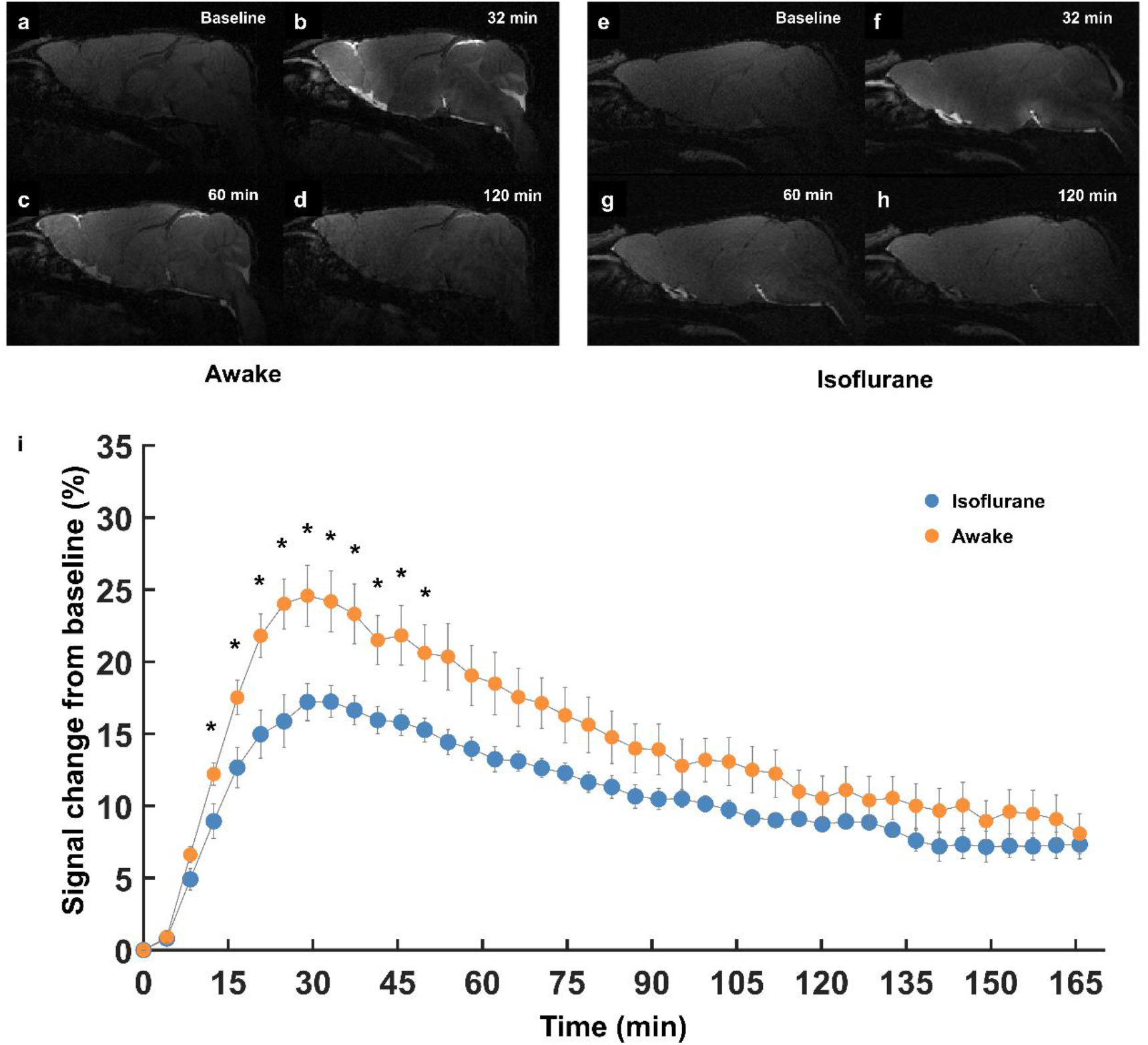
Isoflurane anesthesia reduced the glymphatic function compared to the awake condition using DCE-MRI. (a to d) Representative T1-weighted MRI images in the awake condition before (a), 32min (b), 60min (c) and 120min (d) after infusing Gd-DTPA. (e to h) Representative T1-weighted MRI images in the isoflurane anesthetized condition. i, average time signal curves (TSC) of Gd-DTPA induced signal change from whole brain. *, p < 0.05. Isoflurane, n = 5 animals; Awake, n = 6 animals.

## Discussion

In summary, we developed HEAP-METRIC phase-contrast MRI method, which allows non-invasive and global mapping of low-velocity CSF flow in the mouse brain for the first time. Utilizing this novel technique, we revealed anesthesia induced reduction of CSF flow velocity, compared to the awake state.

To accelerate PC-MRI acquisition, we developed HEAP-METRIC acceleration approach using Fourier basis vectors constructed complex orthogonal matrices to achieve arbitrary number simultaneous slice scanning. This generalized Hadamard encoding scheme achieved 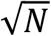 SNR per time improvement, which is critical for global mapping of slow CSF flow in the mouse brain. For previous Hadamard MB encoding method the simultaneously slice number was limited to 2 or 4k (k is an integer) which is mathematically a necessary but not sufficient condition for the construction of a real Hadamard matrix. Our HEAP-METRIC approach uses complex orthogonal matrix to achieve a general form of Hadamard encoding, which is supported by the fact that MRI signal is complex. Thus, any arbitrary number of slices for Hadamard encoding acquisition can be achieved. In practical applications, the maximum MB factor is mainly limited by two hardware factors: minimum dwell time of RF transmitter and maximum output power of RF amplifier. Under those two constrains, we estimate maximum MB factor can reach ~100 in our Bruker scanner for a regular mouse experiment under the conditions: 1000 Watts power of RF amplifier, 3 ms RF duration and 15° flip angle. Furthermore, this new HEAP-METRIC scheme may be extended to other short TR MRI applications, including TOF sequence for vascular imaging, FLASH or SSFP sequence for multi-slice cardiac imaging, and EPI sequence for fMRI, in all of which SNR per time may be increased substantially.

The current multi-band PC-MRI is developed for mapping of low velocity (10^2^ μm/s) flow, including but not limited to mouse CSF flow. CSF plays an important role in the homeostasis and waste clearance in the brain^21,22^. In addition to its mechanical cushion function, CSF also communicates with ISF and maintains the extracellular environment homeostasis^23^, and CSF dynamics perturbation induces cerebral pathologic changes, such as hydrocephalus^24^. Thus, CSF flow dynamics may serve as biomarker for those pathological conditions^25^. In the past decades, various methods were developed to measure the production and absorption of CSF in animal models^26–28^. These approaches, while being highly valuable, are invasive. The nature of the CSF circulation dictates that invasive measurement would likely perturb its dynamics. Therefore, noninvasive methods such as MRI may be complimentary to existing invasive methods. While PC-MRI has been widely used in human for (mostly local) CSF flow measurement^29–31^, global mapping of much lower velocity CSF in rodent is only achieved in the current study with our newly developed HEAP-METRIC scheme (Fig. 1).

With mouse being the predominant animal model in neuroscience research, the current method paves ways for dissecting roles of various genetic, physiological or pathological factors on CSF flow dynamics, and may have further implications on the glymphatic system. It is known that CSF flow is pulsatile and is dependent upon cerebral arterial pulsation^32^, which is considered as the main driver of glymphatic system^33,34^. Therefore, we speculate there might be underlying relationship between CSF flow and glymphatic function. The current study already provided a good example, showing isoflurane anesthesia both significantly reduced CSF velocity and glymphatic function (Fig. 6), compared to the awake state. The exact relationship between CSF flow and glymphatic function will require further examination, possibly utilizing the current HEAP-METRIC PC MRI method.

In conclusion, we developed the novel HEAP-METRIC phase-contrast MRI technique, which enables non-invasive and global mapping of mouse CSF flow velocity. Utilizing this novel technique, we further revealed isoflurane anesthesia induced CSF flow reduction, compared to the awake condition. The technique can be potentially applied in mapping other low velocity flow, and the HEAP-METRIC scheme can also be extended in other MR techniques to significantly boost SNR per time.

## Acknowledgement

This work was supported by the Strategic Priority Research Program of the Chinese Academy of Sciences (Grant No. XDB32030100 to Z. L.), the Shanghai Municipal Science and Technology Major Project (Grant No. 2018SHZDZX05 to Z. L.), the General Program of National Natural Science Foundation of China (Grant No. 81771821 to Z. L.), CAS Pioneer Hundred Talents Program (to Z. L.), the Clinical Research Project of Health Industry of Shanghai Health Committee (Grant No. 20194Y0087 to X. Z.)

## Methods

### 1. Imaging Theory

#### 1.1 Hadamard Encoding APproach of Multi-band Excitation for short TR Imaging aCcelerating (HEAP-METRIC)

A multi-band excitation RF waveform can be expressed by a sum of individual RF pulses with different on-resonance frequencies:

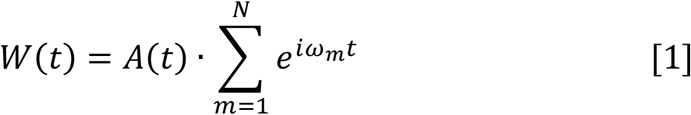

Where W(t) denotes time dependent multi-band RF waveform; A(t) denotes a standard single band selective excitation pulse waveform (e.g., a sinc or hyperbolic secant); N is number of simultaneous bands (or slices); m is the index of band and ω_m_ is the on-resonance frequency of m^th^ band. With Hadamard Encoding phase modulation, a set of N RF waveforms are generated, in which case, Eq. [1] becomes:

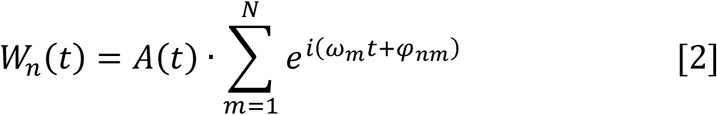

Where n denotes the index of HEAP-METRIC RF pulse; φ_nm_ denotes the Hadamard encoding phase of m^th^ band of n^th^ RF waveform. With elements *e^iφnm^*, an N×N Hadamard Matrix is constructed to be:

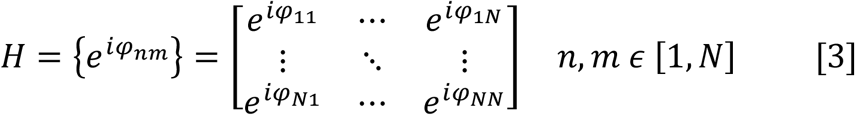

H is orthogonal and satisfies Eq. [4]:

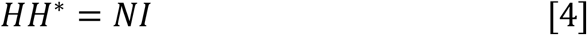

Where I is the identity matrix; H* is the conjugate transpose of H; note that H* = H^T^ when H is real. Previous work^1–4^ all used real Hadamard matrix with all ±1 entries (equivalent to φ_nm_ = 0°or 180°either) thus N is limited to 2 or 4k (k is an integer). Here we generalize Hadamard matrix to be complex thus N can be any arbitrary integer. Fourier basis vectors are used to construct our new Hadamard matrix 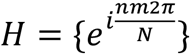.

The MR signal exited by the n^th^ HEAP-METRIC RF pulse is then formulated as:

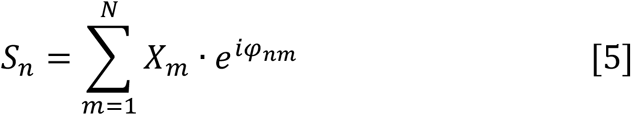

which can also be simplified as Eq. [6]:

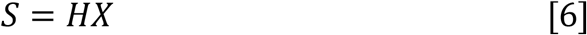

Because the H is orthogonal then each individual band signal X_m_ can be obtained by Eq. [7]:

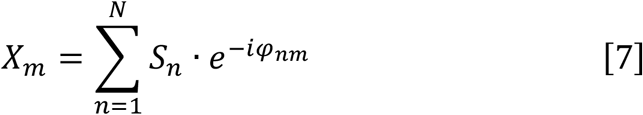

which again can be simplified as Eq. [8]

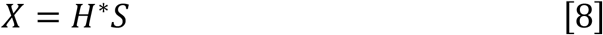

For practical MRI applications, Hadamard decoding by Eq. [7] or [8] can be done either before or after Fourier transform of K space data. N separated scans are required to obtain all N bands images. Comparing with single band RF approach acquiring signal slice by slice, HEAP-METRIC provides N times signal averages with N simultaneous slices with identical acquisition time. Therefore, HEAP-METRIC accelerates image acquisition by a factor of N, thus increases the SNR per time by a factor √N.

#### 1.2 Quantitative CSF Velocity Mapping

The aforementioned HEAP-METRIC method was incorporated into conventional PC-MRI, to substantially speed up acquisition. For in vivo imaging we designed a group of 18 Hadamard encoded eighteen-bands MB RF waveforms and each RF waveform consists of 10000 phase modulated complex points. A Hanning windowed three-lobes-sinc function was used as the base RF waveform A(t).

All excited bands of MB RF pulses were designed to be equally spaced. Adjacent bands were gapped by one band distance thus it needs twice slice-selection (2 transmitting frequencies) to complete a whole brain scan. 36 slices data were finally reconstructed in total. Our HEAP-METRIC PC pulse sequence was implemented on Bruker ParaVision 6.0.1 system. The specific looping structure of a running sequence is as:

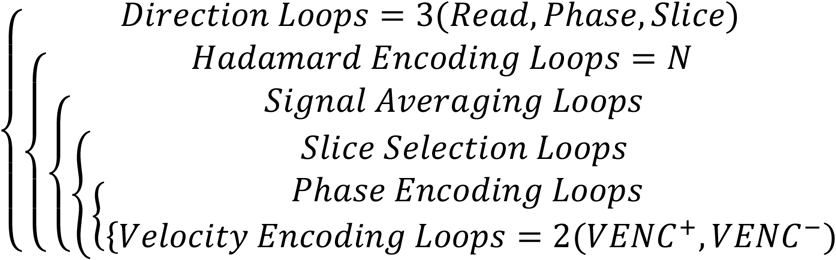

To cancel eddy current effects, we post-processed phase contrast images by spatial polynomial regressing on static tissue data that required a manually drawn global mask excluding flowing substance (e.g., blood or CSF). Results were then subtracted by estimated eddy field^5–8^ . The fitting and correcting procedures were done respectively on data of three different velocity encoding directions (read, phase and slice). Quantitative velocity magnitude and RGB color vector maps were then generated by combining of three directional results. All velocity images were reconstructed and post-processed by our home-built software which was programmed with Microsoft Visual Studio C ++ IDE.

The HEAP-METRIC protocol included in the current study can be openly accessed (https://github.com/ZhifengLiangLab/HEAP-METRIC.git).

### 2. Phantom HEAP-METRIC PC-MRI Experiment

Phantom MRI experiment was carried out to simulate the slow flow (~10^2^ μm/s) and evaluate the sensitivity and accuracy of HEAP-METRIC PC-MRI. A silicone tubing (ID = 2 mm, OD = 3.5 mm, length = 5 m, McMaster-Carr, Douglasville, GA) and a 10 mL syringe (BD, Franklin Lakes, NJ) were integrated into a semi-closed loop fluid flow system using a syringe pump (Model 33, Harvard Apparatus, Holliston, MA). Artificial cerebrospinal fluid (aCSF) was pumped at a control flow rate during MR imaging. The imaging part of the holder was filled with 1% agarose gel and the silicone tubing was placed in the agarose (Fig. S2). This flow phantom system was tested outside the magnet by weighing the aCSF delivered at the set flow rate which resulted in errors less than 10%.

During the phantom experiment, the flow rate was sequentially set to 0.314 μl/s, 0.628 μl/s, 0.942 μl/s, 1.256 μl/s, 1.57 μl/s, 1.884 μl/s, respectively corresponding to 100 μm/s, 200 μm/s, 300 μm/s, 400 μm/s, 500 μm/s, 600 μm/s. The imaging parameters were: TR/TE = 30/9.1 ms, Flip angle = 10°, receiver bandwidth = 100 KHz, FOV = 16×16 mm^2^, matrix size = 180×200, slice thickness = 0.4 mm, VENC value = 0.15 cm/s, pulse duration = 1 ms.

### 3. Animal HEAP-METRIC PC-MRI Experiment

Male adult C57BL/6 mice were used in the current study (14 weeks of age, weighted between 25 and 30 g) with food and water ad libitum. Overall, six mice were imaged during isoflurane anesthesia and seven mice were imaged in the awake condition. All animal experiments were approved by the Animal Care and Use Committee of the Center for Excellence in Brain Science and Intelligence Technology, Chinese Academy of Sciences, Shanghai, China.

For awake imaging group, awake mouse preparation and imaging setup was done according to our previous study^9,10^. Briefly, a head holder was implanted above the animal’s skull for head fixation during imaging. After recovery from the implantation surgery, mice were acclimated to the MRI environment with head fixation and recorded imaging noises.

For anesthetized imaging group, mice were initially induced with 5% isoflurane and endotracheally intubated. During imaging, mice were ventilated and anesthesia was maintained at 1.3% isoflurane delivered by a mixture of oxygen and air (20%: 80%).

All imaging data were obtained on a Bruker 9.4T scanner. Scanning parameters were optimized as: TR/TE = 30/9.1 ms, Flip angle = 10°, receiver bandwidth = 100 KHz, FOV = 16×16 mm^2^, matrix size = 200×200, slice thickness = 0.4 mm, VENC value = 0.15 cm/s, pulse duration = 1 ms. Total scanning time was 21.8 minutes.

### 4. Animal DCE-MRI Experiment

For the study of the glymphatic system, we applied the dynamic contrast enhanced MRI (DCE-MRI) to observe the function of glymphatic system in different conditions. All the included mice referred to the above, and totally five mice were imaged during isoflurane anesthesia and six mice were imaged in the awake condition.

For all imaging groups, mouse preparation and imaging setup were done according to the previous studies^11,12^. Briefly, anesthetized mice were placed in three-point stereotaxic apparatus. A custom-made borosilicate capillary (tip diameter of approximately 100-150 μm) attached to PE-10 tubing filled with aCSF was inserted into the cisterna magna as described previously ^13^. Afterwards, a head holder was implanted above the animal’s skull for head fixation during imaging. After recovery from the implantation surgery, awake mice were acclimated to the MRI environment with head fixation and recorded imaging noises. For the anesthetized imaging group, mice were intubated and ventilated as above during imaging.

All imaging data were obtained on a Bruker 9.4T scanner. T1-weighted 3D-FLASH images were acquired every 4 min. The first three images were baseline images. The paramagnetic contrast agent gadopentetic acid (Gd-DTPA, 938Da) (Byer, German) was infused at the beginning of the fourth scan. The contrast (Gd-DTPA : aCSF = 1 : 40) was infused at a rate of 0.8 μl/min for a total volume of 10μl. 40 scans were acquired continuously for at least 160 min for each study.

### 5. Data Analysis

All data were processed using custom scripts in MATLAB 2018b (MathWorks, Natick, MA), SPM12 (http://www.fil.ion.ucl.ac.uk/spm/) and ITK-SNAP (http://www.itksnap.org/).

First, MR images were normalized to the mouse brain template (https://www.nitrc.org/projects/tpm_mouse) using nonlinear Symmetric Normalization (SyN) algorithm in ANTs. Then we applied the deformation matrix to transform the corresponding flow velocity images into the template space. For the MR image segmentation, a probabilistic atlas (https://www.nitrc.org/projects/tpm_mouse) was aligned with the MRI image and used as a spatial prior for segmentation. Then the MR images were segmented into 5 tissue classes (gray matter, white matter, CSF, tissue outside the brain, and air) using SPM.

To compare global CSF velocities between the awake and anesthetized states, the CSF mask was created by thresholding CSF tissue probability map with probability > 0.9 and with further minor manual correction. The modified CSF mask was applied in the corresponding velocity image to calculate the mean CSF velocity. Allen atlas (https://atlas.brain-map.org/) was utilized to further extract mouse ventricles (lateral ventricle, third ventricle, fourth ventricle and cerebral aqueduct). The above ventricle specific masks were used to obtain mean CSF velocities of each individual ventricle.

For the DCE-MRI data, all images were converted to .nifti format, corrected for head motion, and spatially smoothed. Then the percentage of signal changes from the baseline were calculated in each voxel, as described previously^12^. For each mouse, the time signal curve (TSC) of the whole brain was calculated using MATLAB.

All statistical analysis was performed using GraphPad Prism 8.0 (https://www.graphpad.com/) or MATLAB 2018b. For comparisons between awake and anesthetized groups, unpaired Student’s t-tests were performed. For the phantom MRI experiment and the physiological parameter’s analysis, the linear least squares regression was used for calculation of correlations between the measured velocity (or global CSF velocity) and the set phantom velocity (or heart rate).

